# Microglial activation results in neuron-type-specific increase in mPFC GABAergic transmission and abnormal behavior in mice

**DOI:** 10.1101/2020.06.13.149906

**Authors:** Binliang Tang, Jinxiang Jiang, Lei Wang, Afzal Misrani, Qingwei Huo, Yuanyuan Han, Cheng Long, Li Yang

## Abstract

Neuroinflammation and synaptic dysfunction are two early symptoms of most neurological diseases. However, the mechanisms underlying microglia-associated neuroinflammation in the regulation of synaptic activity remain obscure. We report here that acute neuroinflammation induced by a single-dose proinflammatory cytokine inducer, lipopolysaccharide (LPS), results in enhanced inhibitory postsynaptic currents (IPSCs) of glutamatergic neurons, upregulated levels of GABA_A_R subunits, glutamine synthetase (GS) and vGAT, and downregulated BDNF and pTrkB levels, due to enhanced activation of microglia in the medial prefrontal cortex (mPFC). Blockage of microglial activation by minocycline ameliorated LPS-induced aberrant mIPSCs and associated aberrant protein expression and behavior. Exogenous application of BDNF prior to LPS challenge also ameliorated LPS-induced abnormal mIPSCs. Thus, this study elucidates a critical role for microglia in the neurobiology of GABAergic synaptic dysfunction induced by neuroinflammation, revealing a novel GABAergic signaling pathway that might be targeted therapeutically to treat neuroinflammation-induced abnormal synaptic activity and associated aberrant behavior.

## Introduction

Microglia are immunocompetent cells in the brain that represent approximately 10% of central nervous system (CNS) cells **(Aguzzi et al., 2013; Kettenmann et al., 2013)**. They play multiple roles in shaping and refining neuronal connectivity during development, and mediate activity-dependent synaptic plasticity and neurogenesis in the fully mature CNS **(Bessis et al., 2007; Marinelli et al., 2019; Sierra et al., 2010; Tay et al., 2017; Wu et al., 2015)**. Although an explosion of new findings has provided insight into the involvement of microglia in CNS disorders **(Cai et al., 2020; Salter and Stevens, 2017)**, an understanding of the role of microglia in synaptic signaling, under both healthy and disease conditions, is still elusive.

Neuroinflammation, which is known to influence synaptic function **(Chung et al., 2015; Di Liberto et al., 2018; Habbas et al., 2015)**, has been proposed as a pathological mechanism and potential therapeutic target in numerous neuronal diseases, such as Alzheimer’s disease **(Litvinchuk et al., 2018; Tzeng et al., 2018)**, depression **(Garate et al., 2013; Kaufmann and Menard, 2018)** and autism **(Matta et al., 2019; Pardo et al., 2005)**. However, a question remains whether targeting neuroinflammation might impede the progress of, or even cure, these disease conditions. Microglia respond to inflammatory signals, releasing inflammatory cytokines as well as neuroprotective factors, such as interleukins, tumor necrosis factor (TNF) **(Kim and Joh, 2006; Wang et al., 2015)** and brain-derived neurotrophic factor (BDNF) **(Gomes et al., 2013; Parkhurst et al., 2013)**, the latter being critically involved in synaptic maturation and plasticity **(Salter and Stevens, 2017; Singhal and Baune, 2017; Tremblay et al., 2011)**. Although inflammation and synaptic dysfunction are two early symptoms of most neurological diseases **(Henstridge et al., 2016; Reus et al., 2015; Wyss-Coray and Mucke, 2002)**, the mechanisms underlying the regulation of neuronal activity by microglia-associated neuroinflammation remain to be determined. Limited studies have focused mostly on the link between glutamatergic synaptic alterations and microglial activation **(Haroon et al., 2017; Pascual et al., 2012; Reus et al., 2018; Yang et al., 2019)**, emphasizing subsequent hippocampal dysfunction **(Bie et al., 2019; Dissing-Olesen et al., 2014; Hailer et al., 2001; Riazi et al., 2015; Roumier et al., 2008)**.

GABAergic interneurons constitute up to 25% of the total number of neurons in the cortex, including the medial prefrontal cortex (mPFC) **(Schmidt-Wilcke et al., 2018)**, a brain region that is critical for cognitive and affective processes and that expresses high levels of GABA receptors **(Caballero and Tseng, 2016; Cho et al., 2015)**. GABAergic signals in the PFC are crucial for the regulation and orchestration of glutamatergic pyramidal neuronal assemblies and function **(Kamigaki and Dan, 2017; Sun et al., 2019; Xu et al., 2019)**. However, despite increasing interest in the role of microglia in the regulation of CNS electrical activity, the mechanisms underlying glial regulation of GABAergic function are currently insufficiently understood **(Chen et al., 2014; Choudary et al., 2005; Lee et al., 2010; Mazaud et al., 2019)**.

To address the knowledge gap, we aimed to elucidate the synaptic mechanisms underlying microglial regulation of mPFC synaptic activity in this study. Microglia were activated by a single-dose injection of the proinflammatory cytokine inducer, lipopolysaccharide (LPS), and electrophysiological, biochemical and behavioral evaluations were performed 2 h after LPS injection. In this way, the cellular and molecular alterations resulting from acute neuroinflammation were determined. We show that microglial activation leads to a significantly increased miniature inhibitory, but not excitatory, response in the mPFC pyramidal neurons, i.e. in the postsynaptic component of inhibitory-excitatory synapses. In addition to revealing a critical role for microglia in the regulation of GABAergic synaptic activity in the mPFC, the present study highlights the GABA signaling pathway as a target for the improvement of mPFC brain function during the early stages of infection.

## Results

### Upregulation of mIPSCs in mPFC pyramidal neurons of LPS mice

To explore the early effects of systemic inflammation on synaptic transmission in the mPFC, we first performed whole-cell patch clamp recordings in acute *in vitro* cortical slices containing the mPFC of mice subjected to intraperitoneal (IP) injection of 0.5 mg/kg LPS (denoted ‘2 h-LPS mice’ in this article) or PBS 2 h (Ctrl) earlier. Both miniature excitatory postsynaptic currents (mEPSCs) and miniature inhibitory postsynaptic currents (mIPSCs) of the same pyramidal neuron, in the presence of TTX to block fast sodium channel opening and subsequent action potentials, were recorded at a holding potential of −60 mV and 0 mV, respectively **(Allene et al., 2012; Wang et al., 2019)**. Under the IR-DIC microscope, pyramidal neurons were identified in brain slices by their typical triangular-shaped soma, which distinguishes them from GABAergic interneurons **(Choy et al., 2018; Ramaswamy and Markram, 2015; Stuart et al., 1993)**. The results showed that both the amplitude and frequency of mIPSCs recorded from pyramidal neurons, i.e. postsynaptic cells of inhibitory-excitatory synapse, increased significantly in 2 h-LPS mice compared to Ctrls (amplitude: Ctrl, 8.57 ± 0.36 pA; LPS, 9.91 ± 0.51 pA; *P*=0.039, frequency: Ctrl, 1.34 ± 0.16 Hz; LPS, 2.44 ± 0.34 Hz; *P*=0.0082) ***(Fig. 1B, E-F)***. In contrast, neither the amplitude nor the frequency of mEPSCs differed between the two groups (amplitude: Ctrl, 7.78 ± 0.23 pA; LPS, 8.46 ± 0.32 pA; *P*=0.096, frequency: Ctrl, 1.70 ± 0.35 Hz; LPS, 1.62 ± 0.23 Hz; *P*=0.433) ***(Fig. 1A, C-D)***. As we aimed to evaluate early defects induced by LPS, the data shown below focus on 2 h-LPS and Ctrl mice unless otherwise indicated.

**Figure 1.**
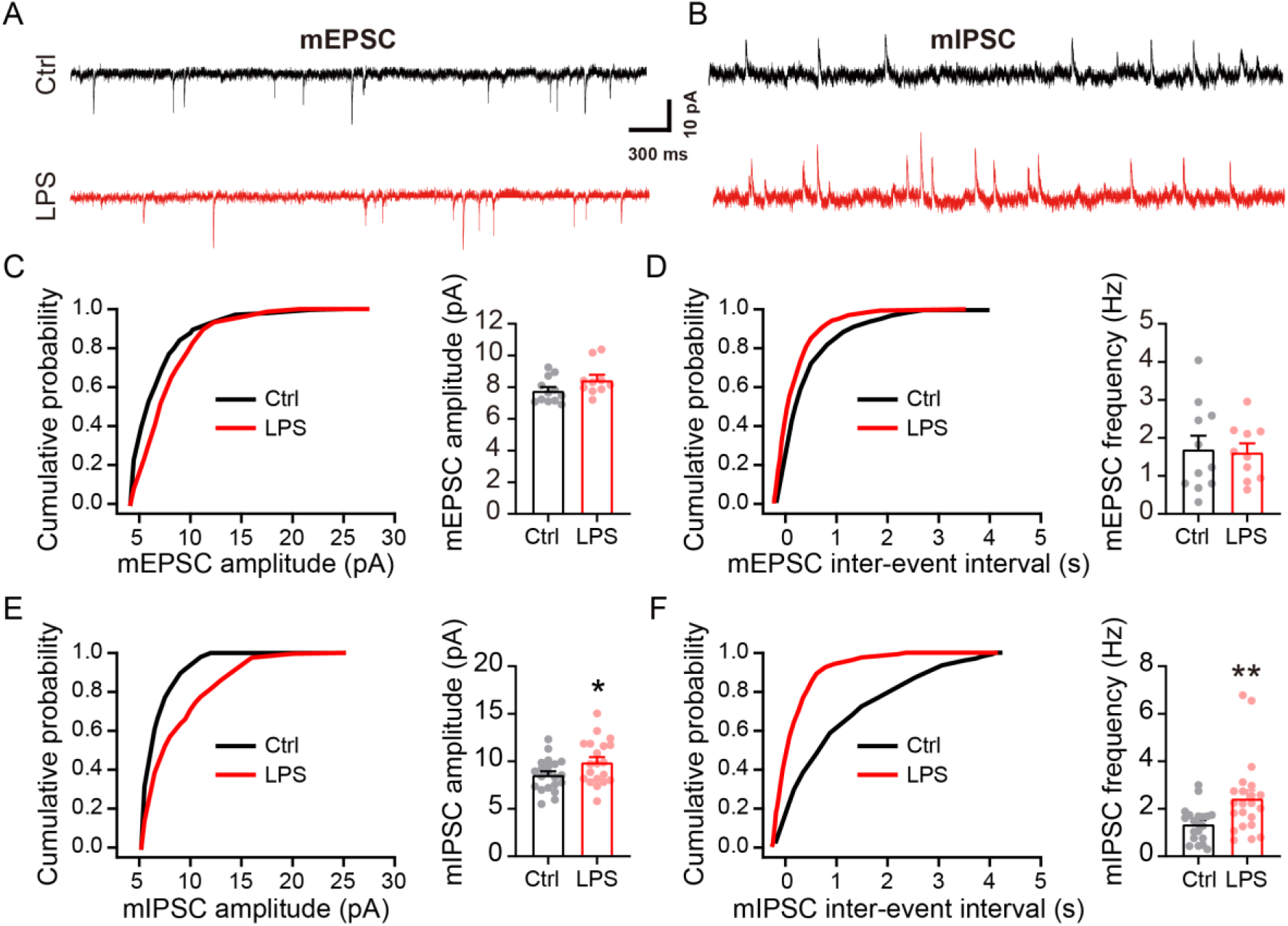
LPS potentiates mIPSC in mPFC pyramidal neurons. Representative traces of mEPSC **(A)** and mIPSC **(B)** recorded from pyramidal neurons of the mPFC. Cumulative distribution and quantification of the mEPSC amplitude **(C)** and frequency **(D)** (Ctrl: n=11 neurons from 3 mice, LPS: n=10 neurons from 3 mice), showing no significant difference in mEPSC amplitude (*P*=0.096) and frequency (*P*=0.433) between Ctrl and LPS mice. Cumulative distribution and quantification of the mIPSC amplitude **(E)** and frequency **(F)** (Ctrl: n=20 neurons from 5 mice, LPS: n=22 neurons from 5 mice). LPS-treated mice showed significantly increased mIPSC amplitude and frequency. Data are presented as means ± SEM. Student’s *t*-test, \**P*<0.05, \*\**P*<0.01.

### Both pre- and postsynaptic mechanisms mediate LPS-induced abnormal inhibition

It is known that mIPSC amplitude correlates with the density and conductance of postsynaptic GABAergic receptors, while mIPSC frequency reflects the number of functional GABA synapses and/or presynaptic probability of release at individual synapses **(Nusser et al., 1997; Nusser et al., 1998; Zucker and Regehr, 2002)**. Thus, the observed increase in mIPSC amplitude and frequency in 2 h-LPS mice suggests that both pre- and postsynaptic activities in mPFC pyramidal neurons are altered. We found further evidence consistent with such alterations when we evaluated IPSCs evoked by single and paired electrical stimulation ***(Fig. 2A)***. mPFC pyramidal neurons in 2h-LPS mice exhibited significantly increased evoked IPSC amplitude (eIPSC) after a single electrical stimulation compared to Ctrls (Ctrl, 204.00 ± 22.92 pA; LPS, 468.22 ± 138.34 pA; *P*=0.031) ***(Fig. 2B)***, indicating that acute administration of LPS modifies postsynaptic activity. We next evaluated the paired-pulse ratio (PPR), a protocol commonly used to evaluate presynaptic function **(Chen et al., 2017; Yang et al., 2009)**, in the mPFC of 2 h-LPS and Ctrl mice. It has been proven that the higher the release probability, the lower the PPR **(Yang et al., 2009; Zucker and Regehr, 2002)**. We found that LPS markedly reduced the PPR of pyramidal neurons (Ctrl, 1.12 ± 0.06; LPS, 0.67 ± 0.06; *P*=0.0002) ***(Fig. 2C)***, revealing an increased probability of presynaptic GABA release in the mPFC **(Regehr, 2012; Zucker and Regehr, 2002)**. Together, these results suggest that acute LPS treatment leads to GABAergic changes both at the pre- and postsynaptic level.

**Figure 2.**
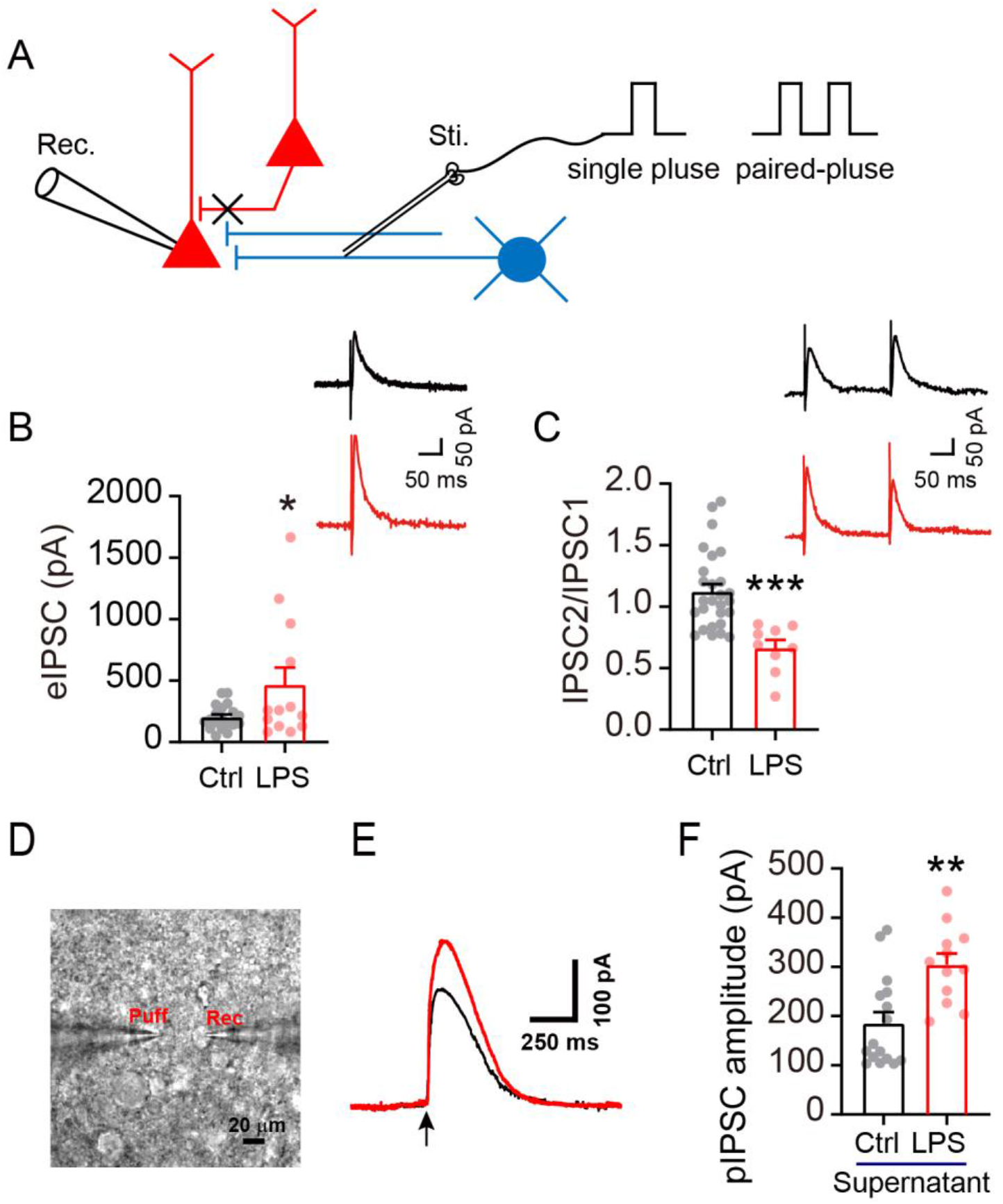
Comparison of eIPSC between pyramidal neurons of Ctrl and LPS mPFC. **(A)** Diagram showing the protocol of electrical stimulation-evoked inhibitory response. Excitatory responses were blocked by CNQX and APV **(B)** Representative traces and quantification showing eIPSC following electrical stimulation (Ctrl: n=19 neurons from 4 mice, LPS: n=13 neurons from 3 mice). **(C)** Representative traces and quantifications of paired-pulse recordings at the 200-ms interval. LPS treatment significantly enhanced the amplitude of eIPSC and meanwhile reduced PPR of eIPSC of pyramidal neurons (Ctrl: n=27 neurons from 5 mice, LPS: n=9 neurons from 3 mice). **(D)** Schematic illustration of the puffing recording paradigm. Recording electrode. Scale, 20 μm. **(E)** Representative traces of pIPSC in respond to puffing-mPFC supernatant extracted from Ctrl and LPS mice, respectively. **(F)** Comparison of pIPSC amplitude between pyramidal neurons of Ctrl (n=16 neurons of 3 mice) and LPS mPFC (n=15 neurons of 3 mice). The pIPSC amplitude was significantly higher in LPS-supernatant compared to Ctrl-supernatant groups, suggesting an increase in GABA content in the mPFC of LPS mice. Data are presented as means ± SEM. Student’s *t*-test, \**P*<0.05, \*\**P*<0.01 and \*\*\**P*<0.001.

### Enhanced GABA content in the mPFC of LPS mice

To further uncover the presynaptic alterations underlying the increased mIPSC frequency in the mPFC of 2-h LPS mice, we conducted a brain-homogenate puff assay to evaluate levels of GABA in the mPFC, as described **(Chen et al., 2017; Feng et al., 2017; Wu et al., 2014)**. Briefly, IPSCs in mPFC pyramidal neurons of normal C57 mice were induced by puffing with mPFC lysates, i.e. the supernatants of mPFC tissue from either 2 h-LPS mice or Ctrls, in whole-cell patch-clamp experiments ***(Fig. 2D)***. In the presence of 20 μM CNQX and 50 μM APV to block excitatory responses **(Chen et al., 2017)**, the amplitude of puff-evoked IPSC (pIPSC) was remarkably elevated in response to LPS-mPFC supernatant compared to that of Ctrls (Ctrl, 185.93 ± 22.98 pA; LPS, 304.21 ± 22.90 pA; *P*=0.001) ***(Fig. 2E-F)***. Given that the postsynaptic cells recorded are normal pyramidal neurons of mPFC, the increase in pIPSC amplitude suggests an increased content of GABA neurotransmitter in the mPFC of LPS mice, which may contribute, at least in part, to the increased mIPSC frequency of pyramidal neurons.

### Unchanged mIPSCs of GABAergic neurons in the mPFC of LPS mice

To evaluate whether acute LPS treatment would affect synaptic transmission of mPFC GABAergic interneurons, which represent about 20-30% of cortical neurons **(Rudy et al., 2011)**, we conducted whole-cell patch recordings in GABAergic interneurons of mPFC using a GAD67^+/GFP^ knock-in mouse line **(Huo et al., 2016; Tamamaki et al., 2003)** in which GABAergic neurons express GFP ***(Fig. 3A)***. Interestingly, LPS changed neither the mEPSC (amplitude: Ctrl, 11.83 ± 0.93 pA; LPS, 10.26 ± 0.45 pA; *P*=0.096, frequency: Ctrl, 2.50 ± 0.38 Hz; LPS, 2.69 ± 0.45 Hz; *P*=0.433) ***(Fig. 3B, D-E)*** nor the mIPSC (amplitude: Ctrl, 11.18 ± 0.55 pA; LPS, 11.58 ± 0.44 pA; *P*=0.565, frequency: Ctrl, 1.75 ± 0.26 Hz; LPS, 1.28 ± 0.25 Hz; *P*=0.357) of GFP-positive interneurons ***(Fig. 3C, F-G)***. These data suggest that LPS potentiated IPSC in the mPFC only when the postsynaptic cells are glutamatergic pyramidal neurons (i.e. at an inhibitory-excitatory synapse) and not in GABAergic interneurons (inhibitory-inhibitory synapse).

**Figure 3.**
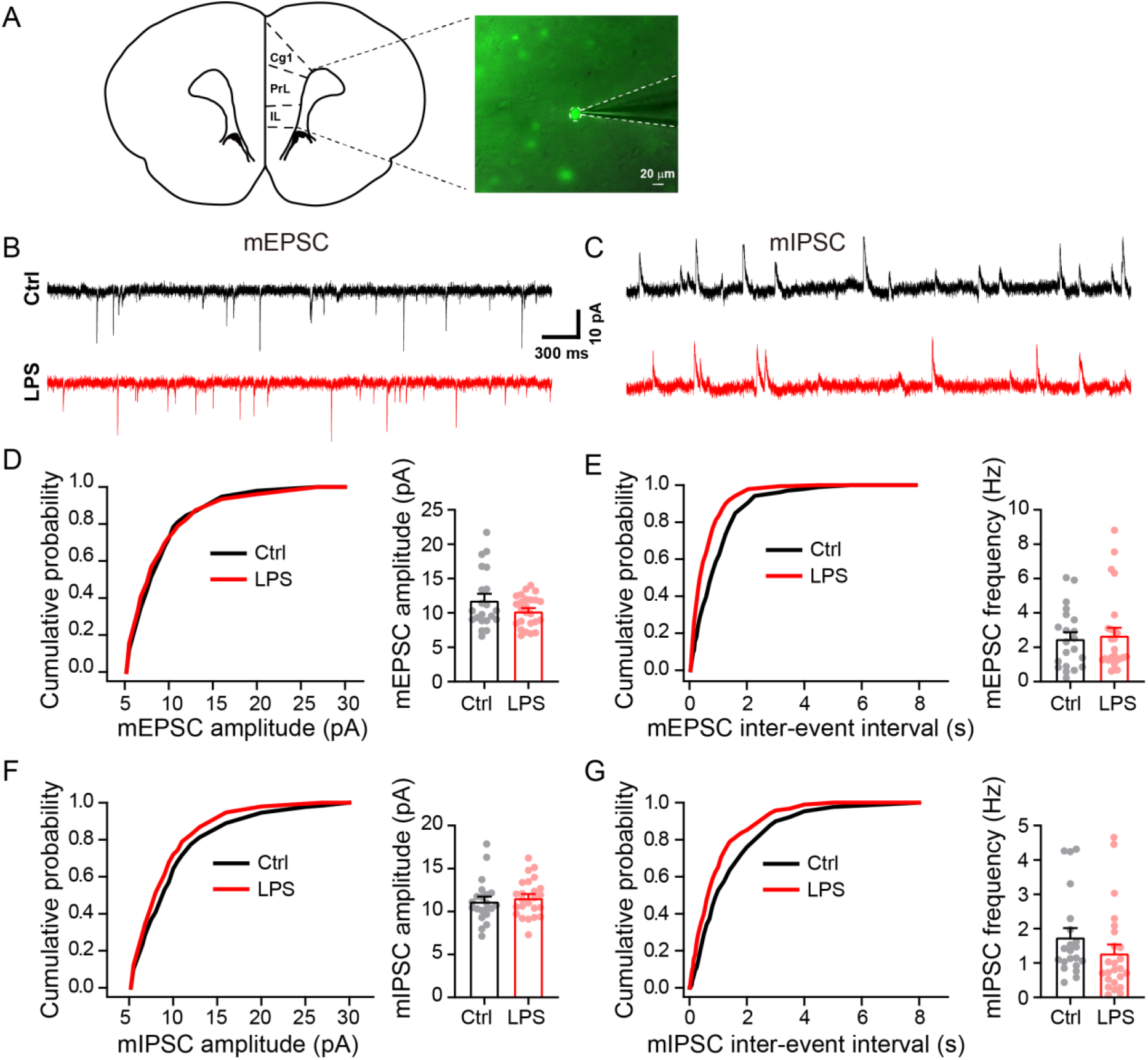
Excitatory and inhibitory synaptic transmission of GAD67-GFP interneurons remained normal in the mPFC of LPS mice. **(A)** Diagram of mPFC (Cg1, cingulate cortex, area 1; PrL, prelimbic cortex; IL, infralimbic cortex) (left) and epifluorescent image of GFP-labeled GABAergic interneurons for the whole-cell recording (right) in the mPFC of GAD67^+/GFP^ mice. Scale, 20 μm. Representative traces of mEPSC **(B)** and mIPSC **(C)** recorded from GABAergic interneurons of LPS and Ctrl groups. Cumulative distribution and quantification of the mEPSC amplitude **(D)** and frequency **(E)** (Ctrl: n=21 neurons from 5 mice, LPS: n=25 neurons from 5 mice), and the mIPSC amplitude **(F)** and frequency **(G)** (Ctrl: n=21 neurons from 5 mice, LPS: n=20 neurons from 5 mice). Both mIPSC and mEPSC remained unaffected in GABAergic neurons after LPS treatment (mEPSC amplitude, *P*=0.096; mEPSC, *P*=0.433; mIPSC amplitude, *P*=0.565; mIPSC frequency, *P*=0.357). Data are presented as means ± SEM. Student’s *t*-test.

### Altered levels of GABA signaling proteins in the mPFC of LPS mice

The α1 and β2 subunits, which are the predominant postsynaptic GABA_A_ receptor subunits in the mature cortex, mediate the vast majority of GABA function in the PFC **(Akbarian et al., 1995; Datta et al., 2015; Hashimoto et al., 2009)**. An increase in IPSC amplitude commonly implies greater involvement and activation of postsynaptic GABA_A_Rs **(Kilman et al., 2002; Luscher et al., 2011)**. In line with the above notion, our western blotting analysis demonstrated significantly enhanced expression of GABA_A_R α1 and β2 proteins, but not GABA_A_R α5, in the mPFC of LPS mice (GABA_A_R α1: *P*=0.0004; β2: *P*=0.01; α5: *P*=1.00) ***(Fig. 4A)***. RT-PCR analysis showed that mRNA levels of GABA_A_R α1 and β2 were also increased (GABA_A_R α1: *P*=0.01; β2: *P*=0.004) ***(Fig. 4B)***, indicating transcriptional regulation of the two subunits by LPS.

**Figure 4.**
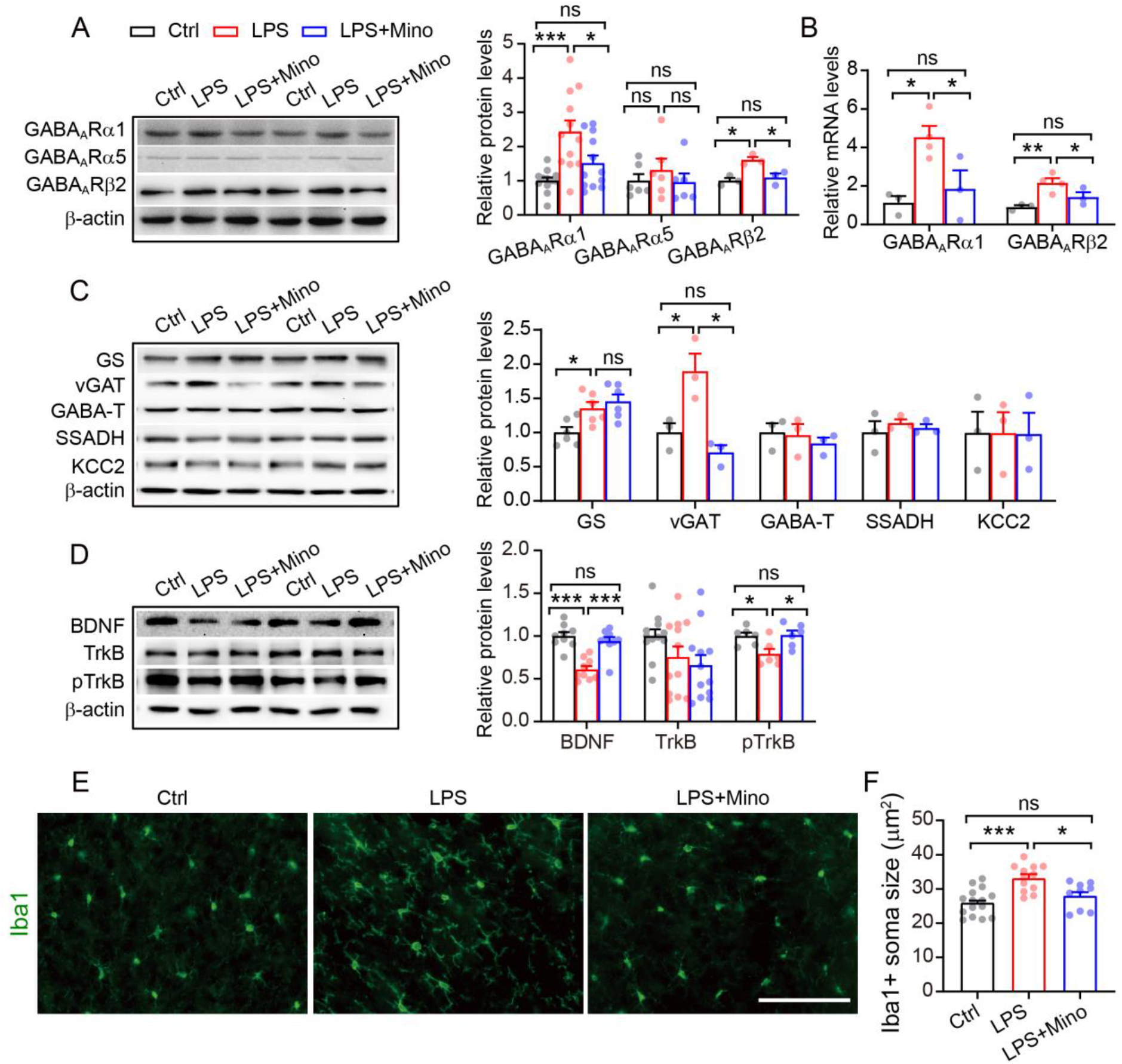
LPS-induced abnormal expression of GABA signaling proteins is dependent on microglia activation. **(A)** Representative immunoblots of mPFC extracts from Ctrl, LPS and LPS+Mino mice (left), and quantification of the immunoblots (right) revealing significantly increased levels of GABA_A_R α1 (n=12 mice/each group) and β2 (n=6 mice/each group,). Altered protein expression was ameliorated by minocycline. **(B)** Relative mRNA levels of GABA_A_R α1 and β2 subunits in the mPFC of Ctrl, LPS and LPS+Mino mice analyzed by real-time PCR and presented as fold change against Ctrl counterpart (Ctrl, n=3; LPS, n=4; LPS+Mino, n=3). **(C)** Representative immunoblots and quantification of the immunoblots reveal significantly increased levels of GS and vGAT which can be ameliorated by minocycline (Ctrl, n=3; LPS, n=3; LPS+Mino, n=3). **(D)** Representative immunoblots and quantification of the immunoblots show significantly decreased levels of BDNF and p-TrkB which can be ameliorated by minocycline (n=6 mice/each group). **(E)** Immunofluorescent staining of Iba1 on the mPFC sections. Scale, 100 μm. **(F)** Iba1^+^ soma size was compared among three groups (Ctrl, n=5; LPS, n=4; LPS+Mino, n=3). Microglia of mPFC became moderately active as shown by larger cell bodies in LPS mice, which could be efficiently suppressed by minocycline (*P*=0.243 against saline-treated control). Data are presented as means ± SEM. One-way ANOVA followed by Bonferroni’s *post hoc,* or Turkey’s multiple comparisons, \**P*<0.05, \*\**P*<0.01, \*\*\**P*<0.001 and ns, not significant.

Glutamine synthesizing enzyme, glutamine synthetase (GS), is used as a precursor for synthesis of GABA in GABAergic neurons **(Hertz, 2013; Rowley et al., 2012)**. Moving GABA synthesized into synaptic vesicles is conducted by a vesicular GABA transporter (vGAT) **(Jin et al., 2003; Jones, 2012)**. LPS challenge has been shown to increase GS levels **(Wang et al., 2018)**. Changes in GS and vGAT expression directly regulate neurotransmitter release **(Gomes et al., 2011; Huo et al., 2017)**. We thus evaluated whether the LPS-invoked increase in mIPSC frequency *via* presynaptic mechanisms involves abnormal expression of GS and vGAT. Indeed, western blotting showed that levels of GS and vGAT increased significantly in 2h-LPS, compared to Ctrl, mPFC (GS: *P*=0.007; vGAT: *P*=0.04) ***(Fig. 4C)***, suggesting possible increase in GABA synthesizing and loading into vesicles, which is in line with increased mIPSC frequency ***(Fig. 1F)*** and pIPSC ***(Fig. 2D-F)***. In comparison, levels of GABA degradation related proteins, such as GABA transaminase (GABA-T) and succinic semialdehyde dehydrogenase (SSADH) **(Parviz et al., 2014; Roth and Draguhn, 2012)**, and potassium-chloride cotransporter 2 (KCC2) which hyperpolarizes GABA_A_R mediated response **(Chen et al., 2017; Jaenisch et al., 2010)** remained unchanged (GABA-T: *P*=1.00; SSADH: *P*=1.00; KCC2: *P*=1.00) ***(Fig. 4C)***. Interestingly, LPS significantly reduced levels of BDNF and its receptor, phosphorylated tropomyosin-related kinase B (pTrkB) (BDNF: *P*=0.0001; pTrkB: *P*=0.039) ***(Fig. 4D)***. It is worth noting that expression of glutamatergic excitation-related proteins, such as NMDAR1 (N1R), NMDAR2B (NR2B) and glutamate receptor 1 (GluR1, AMPAR subtype), remained similar between the two groups (NR1: *P*=0.74; NR2B: *P*=0.30; GluR1: *P*=0.93) ***(Suppl. Fig. 1)***.

### Minocycline prevents LPS-induced abnormal protein expression in the mPFC

A single dose of peripheral LPS is sufficient to activate microglia and induces neuroinflammation in the CNS **(Murray et al., 2011; Pascual et al., 2012; Qin et al., 2007)**. To determine whether LPS-induced alterations resulted from microglial activation, we compared protein levels of GABA_A_R subunits in LPS-treated mice with or without minocycline pretreatment. Minocycline is a specific inhibitor of microglial activation **(Huang et al., 2014; Zink et al., 2005)**, which attenuates LPS-induced neuroinflammation **(Henry et al., 2008)**. To mitigate the possible side effects of minocycline, such as dizziness, nausea and severe headache **(American Society of Health System Pharmacist, 2019)**, which would affect brain activities, minocycline was injected prior to LPS challenge. Specifically, mice were pretreated with either minocycline (50 mg/kg, IP) or PBS once daily for three consecutive days followed by LPS injection on the third day. Brain tissues were obtained 2 h after IP injection of LPS. Western blotting demonstrated that minocycline pretreatment caused a significant normalization in protein levels of the α1 and β2 subunits (GABA_A_R α1: *P*=0.03; β2: *P*=0.03) ***(Fig. 4A)***, in addition to levels of mRNA (GABA_A_R α1: *P*=0.03; β2: *P*=0.02) ***(Fig. 4B)***, in the mPFC of LPS mice. Minocycline also ameliorated expression of vGAT, BDNF and pTrkB (vGAT: *P*=0.01; BDNF: *P*=0.0001; pTrkB: *P*=0.03) ***(Fig. 4C-D)***. These results show that LPS-induced microglial activation contributes to abnormal expression of the above proteins and subsequent alterations in GABAergic synaptic transmission.

### LPS induces microglial activation in the mPFC

To causally prove the contribution of microglia activation to the alteration of GABAergic inhibition and related proteins identified in 2h-LPS mice, we next examined whether 2 h-LPS treatment indeed activates microglia in the mPFC. Utilizing immunofluorescent staining with anti-Iba1 antibody, which is widely used to detect microglia under both normal and pathological conditions in mice **(Sasaki et al., 2001)**, we observed a remarkably enhanced Iba1 signal in the mPFC of LPS mice (*P*=0.0001) ***(Fig. 4E-F)***. Moreover, pretreating the mice for three days with minocycline prevented LPS-induced microglial activation (*P*=0.02) ***(Fig. 4E-F)***. Although activated microglia are known to trigger the induction of astrocytes **(Liddelow et al., 2017)**, no reactive astrocytes, as evaluated by GFAP staining, were observed in the mPFC of either Ctrl or LPS mice ***(Suppl Fig. 2)***, probably because the dose and duration of the LPS insult was not sufficient to activate astrocytes in this brain area. Together, these studies suggest a pivotal role of microglia in the transcriptional regulation of GABAergic receptors in acutely infected mPFC.

### Minocycline ameliorates aberrant mIPSC in mPFC pyramidal neurons

GABA_A_RS determine the amplitude of IPSC **(Nusser et al., 1997; Nusser et al., 1998)**. Because it abrogates the upregulation of GABA_A_R subunits, minocycline might also affect mIPSC amplitude in the mPFC pyramidal neurons of LPS-treated mice. We therefore subjected mice to the same minocycline injection and LPS challenge described above. In line with the reduction of GABA_A_R α1 and β2 subunit levels by minocycline, mIPSC amplitude was significantly reduced in the mPFC of LPS mice pretreated with minocycline compared to LPS alone (LPS, 13.64 ± 0.51 pA; LPS+Mino, 11.43 ±0.49 pA, *P*=0.02) ***(Fig. 5B, C)***. In contrast, minocycline showed no effect on the amplitude of Ctrls (Ctrl, 10.78 ± 0.70 pA;

**Figure 5.**
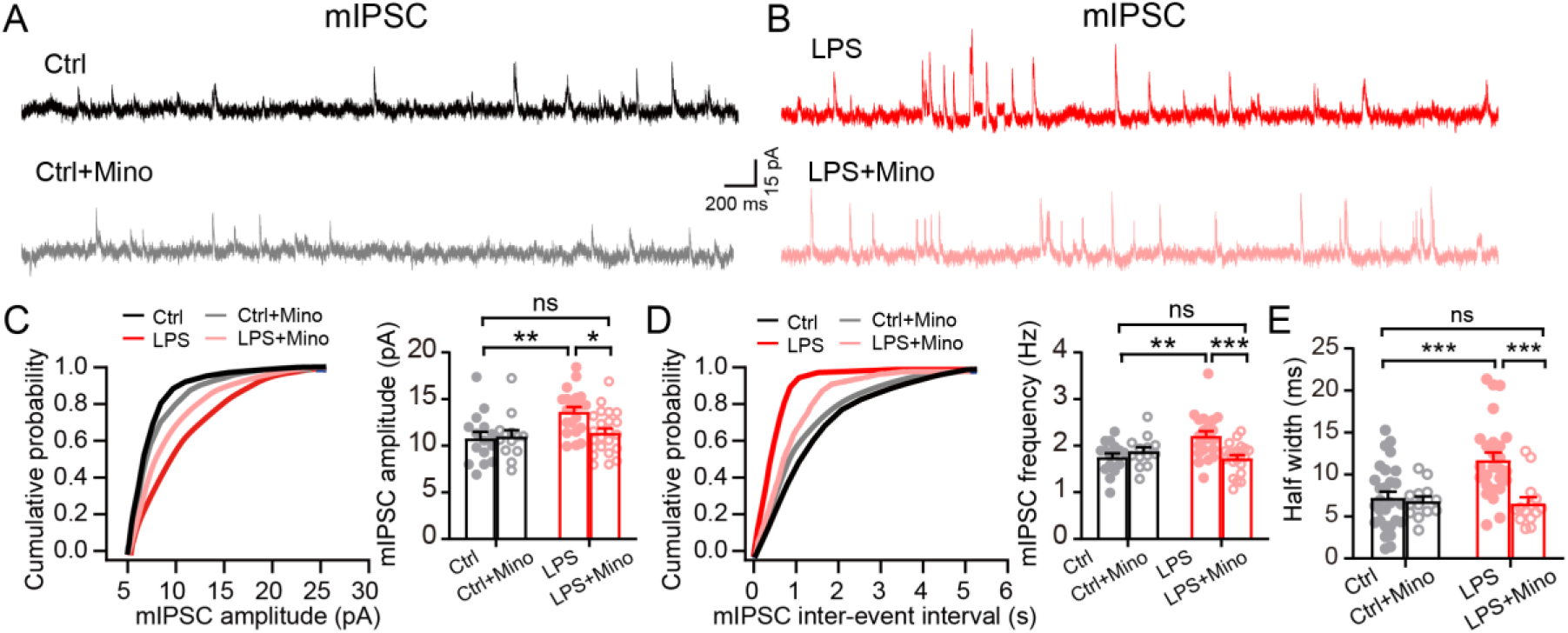
Minocycline ameliorates LPS-induced abnormal mIPSC. Representative traces of mIPSC of pyramidal neurons in the presence and absence of minocycline pretreatment in the Ctrl **(A)** and LPS mice **(B)**. Cumulative distribution and quantification of the mIPSC amplitude **(C)** and frequency **(D)** in the presence and absence of minocycline. Minocycline ameliorated mIPSC amplitude and frequency in LPS mice (LPS: n=21 neurons from 4 mice, LPS+Mino: n=22 neurons from 5 mice), without affecting mIPSC amplitude (*P*=1.00) and frequency (*P*=1.00) in Ctrls (Ctrl: n=16 neurons from 4 mice, Ctrl+Mino: n=15 neurons from 4 mice). **(E)** Significantly increased half-width of mIPSC in LPS mice, and normalization of half-width between Ctrl and LPS mice by minocycline. Data are presented as means ± SEM. Two-way ANOVA followed by Bonferroni’s *post hoc* test, \**P*<0.05, \*\**P*<0.01 and \*\*\**P*<0.001 and ns, not significant.

Ctrl+Mino, 10.98 ± 0.68 pA; *P*=1.00) ***(Fig. 5A, C)***. Moreover, minocycline normalized the amplitude difference between Ctrl and LPS (*P*=0.005 between Ctrl and LPS; and *P*=1.00 between Ctrl and LPS+Mino) ***(Fig. 5C)***. Interestingly, minocycline also ameliorated the mIPSC frequency difference between mPFC of LPS- and LPS/minocycline-mice (LPS, 2.21 ± 0.10 Hz; LPS+Mino, 1.72 ± 0.07 Hz; *P*=0.0007) with no effect on the frequency of Ctrls (Ctrl, 1.75±0.08 Hz; Ctrl+Mino, 1.87±0.09 Hz; *P*=1.00) ***(Fig. 5D)***. The frequency difference between Ctrl and LPS animals was also normalized (*P*=0.004 between Ctrl and LPS, and *P*=1.00 between Ctrl and LPS+Mino) ***(Fig. 5D)***. Additionally, acute LPS treatment significantly increased mIPSC half-width, which is an electrophysiological feature representing an immature stage **(Ali et al., 2000; Mitoh et al., 2008)**. Moreover, pretreatment with minocycline also ameliorated half-width (Ctrl, 7.23 ±0.73 ms; LPS, 11.71 ±0.89 ms; Ctrl+Mino, 6.82 ±0.58 ms; LPS+Mino, 6.55 ±0.73 ms; *P*=0.0002 between Ctrl and LPS, *P*=1.00 between Ctrl and LPS+Mino, *P*=0.0004 between LPS and LPS+Mino) ***(Fig. 5E)***. In the analysis in amplitude, frequency and half-width of mIPSC, the statistics showed significant interactions between treatment and absence/presence of minocycline injection (amplitude: *P*=0.046, frequency: *P*=0.002, half-width: *P*=0.008). Together, the results of the present study indicate that LPS potentiation of mPFC inhibitory synaptic activities in pyramidal neurons depends on LPS-induced microglial activation.

### Probing the molecular pathway underlying microglial activation-induced aberrant GABAergic transmission

Having observed that activation of microglia results in the upregulation of GABAergic synaptic function, we next asked how activated microglia affect GABAergic activities in the mPFC pyramidal neurons of LPS mice. Microglia have long been considered a potential source of BDNF **(Blank et al., 2014; Coull et al., 2005; Marinelli et al., 2019; Parkhurst et al., 2013)**, which attenuates GABAergic transmission in hippocampus **(Tanaka et al., 1997)**. Given that we found LPS treatment to reduce BDNF levels in the current study, which is consistent with another report **(Golia et al., 2019)**, we reasoned that the enhanced IPSC in the mPFC of 2 h-LPS mice may be a consequence of BDNF downregulation. If this is the case, treatment of LPS-challenged mPFC with BDNF should prevent the LPS-induced increase in GABAergic synaptic transmission. Accordingly, we first evaluated the dose-response effect of LPS on the mIPSCs of mPFC pyramidal neurons. Brain slices in the recording chamber were perfused with normal aCSF (i.e. without LPS; 0 ng/ml), or with aCSF containing LPS at a final concentration of 50 ng/ml, 100 ng/ml or 200 ng/ml ***(Fig. 6A)***. LPS incubation, at all doses tested, significantly increased both the mIPSC amplitude (0 ng/ml-Ctrl, 10.69 ± 0.34 pA, 50 ng/ml, 12.48 ± 0.40 pA; 100 ng/ml, 12.73 ± 0.50 pA, 200 ng/ml, 12.65 ± 0.48 pA; *P*=0.0319, 0.0097, 0.0142, respectively) and frequency (0 ng/ml-Ctrl, 1.74 ± 0.10 Hz, 50 ng/ml, 2.14 ± 0.10 Hz; 100 ng/ml, 2.30 ± 0.11 Hz, 200 ng/ml, 2.14 ± 0.09 Hz; *P*=0.0498, 0.0016, 0.0436, respectively) ***(Fig. 6B-C)***. Next, we compared the effect of BDNF on the mIPSCs of mPFC slices treated with 50 ng/ml LPS. The mIPSCs were recorded after preincubation of the slice with or without BDNF-containing aCSF for 20 min followed by perfusion with aCSF containing 0 (Ctrl) or 50 ng/ml LPS for 10 min ***(Fig. 6D-E)***. The results demonstrated that preincubation with BDNF at 20 ng/ml significantly reduced the amplitude of mIPSC following LPS treatment (LPS+BDNF (0 ng/ml), 14.32 ± 0.51 pA; LPS+BDNF (20 ng/ml), 11.33 ± 0.56 pA; *P*=0.0007). But a higher dose (50 ng/ml) was necessary to abrogate the increase in both mIPSC amplitude (LPS+BDNF (0 ng/ml), 14.32 ± 0.51 pA; LPS+BDNF (50 ng/ml), 10.80 ± 0.41 pA; *P*=0.000003) and frequency (LPS+BDNF (0 ng/ml), 2.15 ± 0.09 Hz; LPS+BDNF (50 ng/ml), 1.54 ± 0.44 Hz; *P*=0.000005) in mPFC pyramidal neurons ***(Fig. 6F-G)***. In contrast, neither 20 ng/ml nor 50 ng/ml affected mIPSC in Ctrls (amplitude: 0 ng/ml-Ctrl, 10.69 ± 0.48 pA; 20 ng/ml, 9.79 ± 0.67 pA; 50 ng/ml, 10.37 ± 0.40 pA; *P*=0.486, frequency: 0 ng/ml-Ctrl, 1.69 ± 0.12 Hz; 20 ng/ml, 1.72 ± 0.14 Hz; 50 ng/ml, 1.62 ± 0.09 Hz; *P*=0.742) ***(Fig. 6F-G)***. Two-way ANOVA analysis displayed that there were remarkably interactions between treatment and different dose of BDNF preincubation in both amplitude and frequency of mIPSC (amplitude: *P*=0.004, frequency: *P*=0.01).

**Figure 6.**
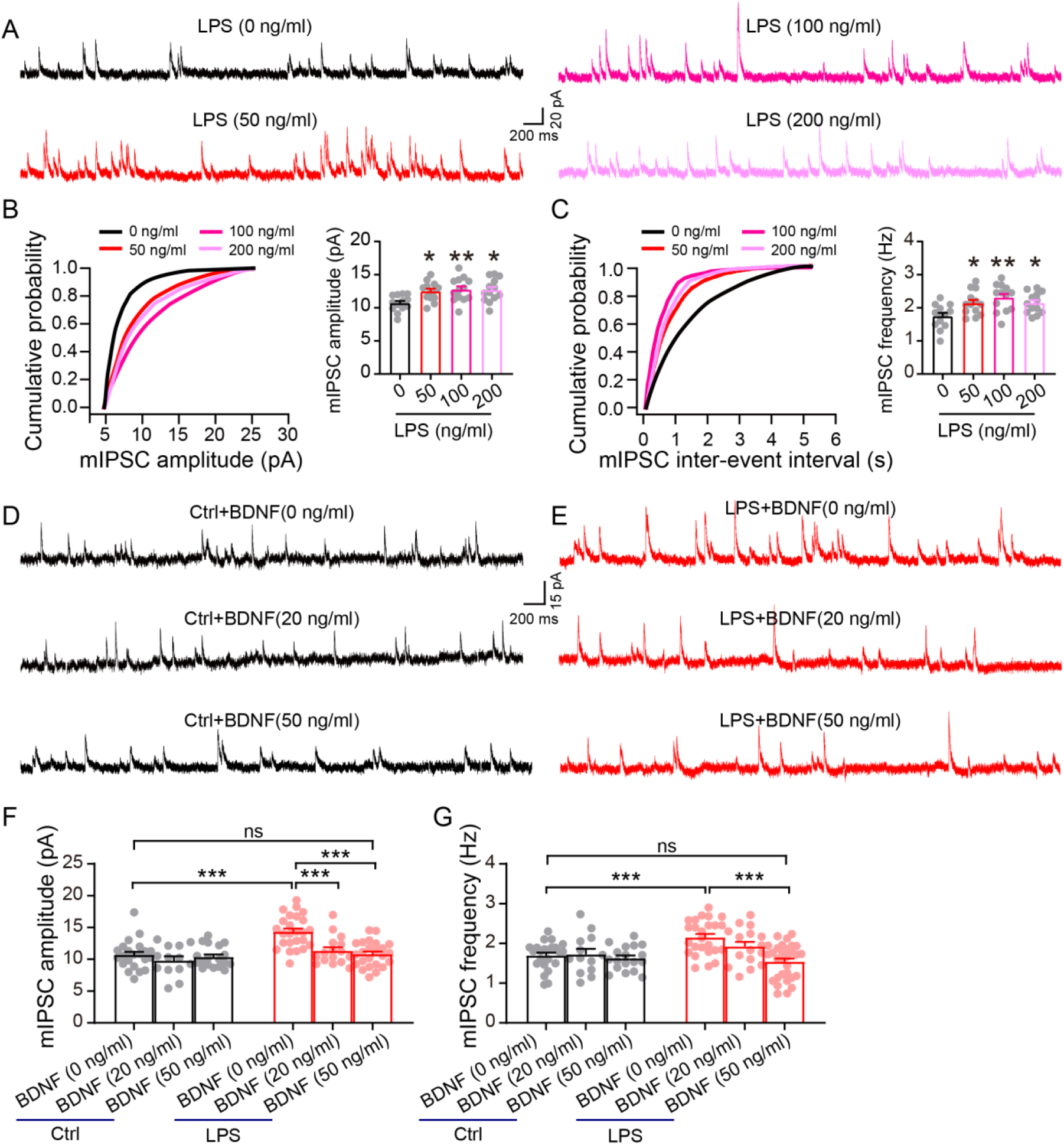
BDNF preincubation prevents LPS-induced mIPSC defects. **(A)** Representative mIPSC traces recorded from acute mPFC slices under graded concentrations of LPS. Cumulative distribution and quantification of the mIPSC amplitude **(B)** and frequency **(C)**. LPS incubation induced significant increases in both mIPSC amplitude and frequency of mPFC pyramidal neurons at all doses tested (n= 13-30 neurons of 4-8 mice). **(D, E)** Representative mIPSC traces recorded from acute mPFC slices under the preincubation of BDNF at the indicated doses (0, 20 and 50 ng/ml) followed by recording in Ctrl **(D)** or in 50-ng/ml LPS treated slices **(E)**. Quantification of the effect of BDNF on mIPSC amplitude **(F)** and frequency **(G)** in Ctrl (n=22 neurons of 7 mice for 0 ng/ml BDNF; n=13 neurons of 4 mice for 20 ng/ml BDNF; and n=18 neurons of 8 mice for 50 mg/ml BDNF) and LPS groups (n=25 neurons of 8 mice for 0 ng/ml BDNF; n=16 neurons of 4 mice for 20 ng/ml BDNF; and n=30 neurons of 8 mice for 50 mg/ml BDNF). BDNF at 20 ng/ml only prevented LPS-induced abnormality of mIPSC amplitude, while 50 ng/ml BDNF prevented both mIPSC amplitude and frequency from LPS-induced abnormalities. Data are presented as means ± SEM. One- or two-way ANOVA followed by Bonferroni’s *post hoc* test, \**P*<0.05, \*\**P*<0.01 and \*\*\**P*<0.001 and ns, not significant.

### Blocking microglial activation attenuates abnormal behavior induced by LPS

LPS has been shown to cause abnormal behavior in mice, such as the suppression of locomotor and exploratory activity, and modifications of social behavior, as early as 2 h after injection **(Dantzer et al., 2008; Duan et al., 2018; Henry et al., 2008)**, perhaps as a strategy for focusing the host’s resources on reducing the spread of infection **(Berg et al., 2004; Bluthe et al., 2000)**. In this study, 2 h-LPS mice demonstrated a significantly reduced total distance and centre time in the OFT ***(Fig. 7A, C-D)***, decreased open arm entries in the EPM ***(Fig. 7B, E)*** and increased immobility in the TST ***(Fig. 7F)***. Importantly, pretreatment with minocycline for three consecutive days attenuated this abnormal behavior ***(Fig. 7A-F)***. Depression-like behavior in mice is usually measured by consumption of sucrose solution, as depressed mice consume less sucrose than normal controls **(Scheggi et al., 2018; Yan et al., 2018)**. We found that sucrose consumption decreased significantly, reaching its lowest level one-day after LPS challenge ***(Fig. 7G)***. Moreover, pretreatment with minocycline for three days before LPS injection significantly increased consumption of sucrose ***(Fig. 7G)***. These results suggest that activated microglia during the early phase of inflammation may have important pathological implications, and blocking microglial activation could prevent, or at least ameliorate, inflammation-induced abnormal behavior.

**Figure 7.**
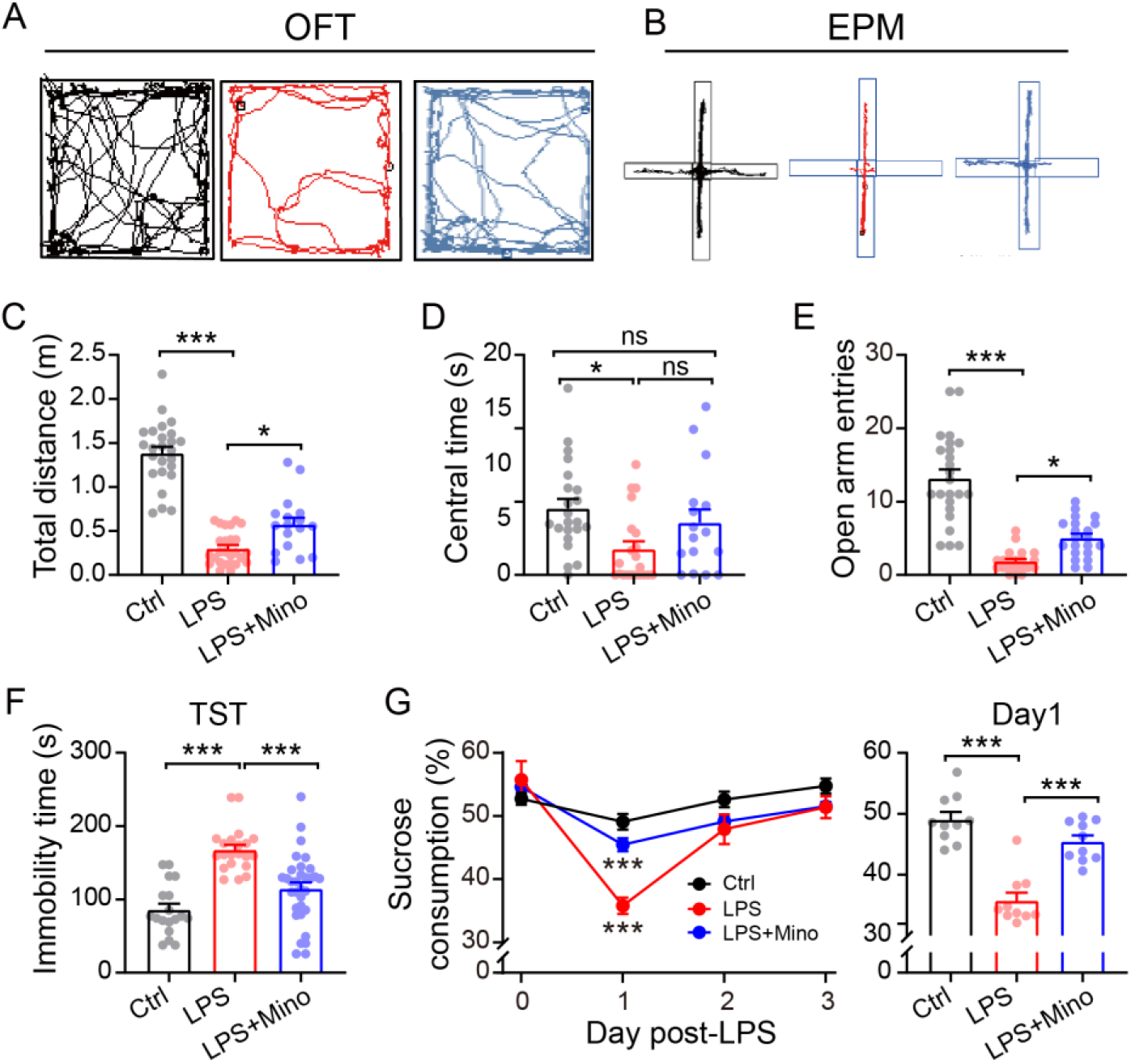
Microglia activation underlies LPS induced abnormal behavior. Representative moving traces of the animals in the OFT **(A)** or the EPM **(B)** during a 5 min exploration period. **(C, D)** LPS mice showed significantly reduced total distance travelled and central time in the OFT. **(E)** The number of open-arm entry reduced significantly in LPS compared to Ctrl in the EPM (Ctrl, n=24; LPS, n=21; LPS+Mino, n=21). **(F)** TST showing significantly increased time spent immobile in LPS mice (Ctrl, n=18; LPS, n=21; LPS+Mino, n=32). **(G)** Significantly reduced sucrose assumption in LPS mice in the SPT (Ctrl, n=10; LPS, n=10; LPS+Mino, n=10). Importantly, pretreatment with minocycline ameliorated above LPS-induced sickness behavior **(C-G)**. Data are presented as means ± SEM. One-way ANOVA followed by Bonferroni’s *post hoc* test, \**P*<0.05, \*\*\**P*<0.001, and ns, not significant.

## Discussion

Here, we show that activation of microglia in the mPFC occurs 2 h after LPS challenge, resulting in increased inhibitory synaptic transmission in the pyramidal neurons of inhibitory-excitatory synapses, upregulated levels of GABAergic signaling proteins and abnormal behavior. Given that mIPSC frequency, GS, vGAT and PPR are presynaptic properties **(Chen et al., 2017; Yang et al., 2009; Zucker and Regehr, 2002)**, while mIPSC amplitude and levels of postsynaptic GABA receptors are postsynaptic properties **(Nusser et al., 1997; Nusser et al., 1998)**, we propose that the altered GABAergic synaptic response in the mPFC pyramidal neurons of 2 h-LPS mice involves changes at both pre- and postsynaptic sites as summarized ***(Fig. 8)***. Specifically, the evidence that LPS induces presynaptic alteration includes 1) an increase in the frequency of mIPSC and a reduction in PPR of eIPSC; 2) upregulated levels of GS and vGAT which likely result in increased GABA accumulation and thus GABA content; and 3) an increase in pIPSC amplitude in mPFC pyramidal neurons of normal C57 mice puffed with mPFC supernatant, sort of presynaptic origin, from LPS mice. That postsynaptic GABAergic changes contribute directly to an altered inhibitory response is supported by 1) an increase in the amplitude of both mIPSC and eIPSC; and 2) upregulation in levels of GABA_A_R α1 and β2 subunits. Interestingly, BDNF/TrkB pathway negatively regulated GABAergic synaptic transmission in LPS-treated mPFC. Moreover, although glutamate (Glu) can be converted to glutamine (Gln) by GS in the astrocytes **(Hertz, 2013; Wang et al., 2018)**, unchanged levels of astrocytic activation in this study suggest a possible increase in microglia metabolization of Glu to Gln *via* GS activity in inflammatory state **(Nakajima et al., 2015)**, which may be crucial to upregulation of GABA amount and subsequent mIPSC frequency in the mPFC of LPS challenged mice.

**Figure 8.**
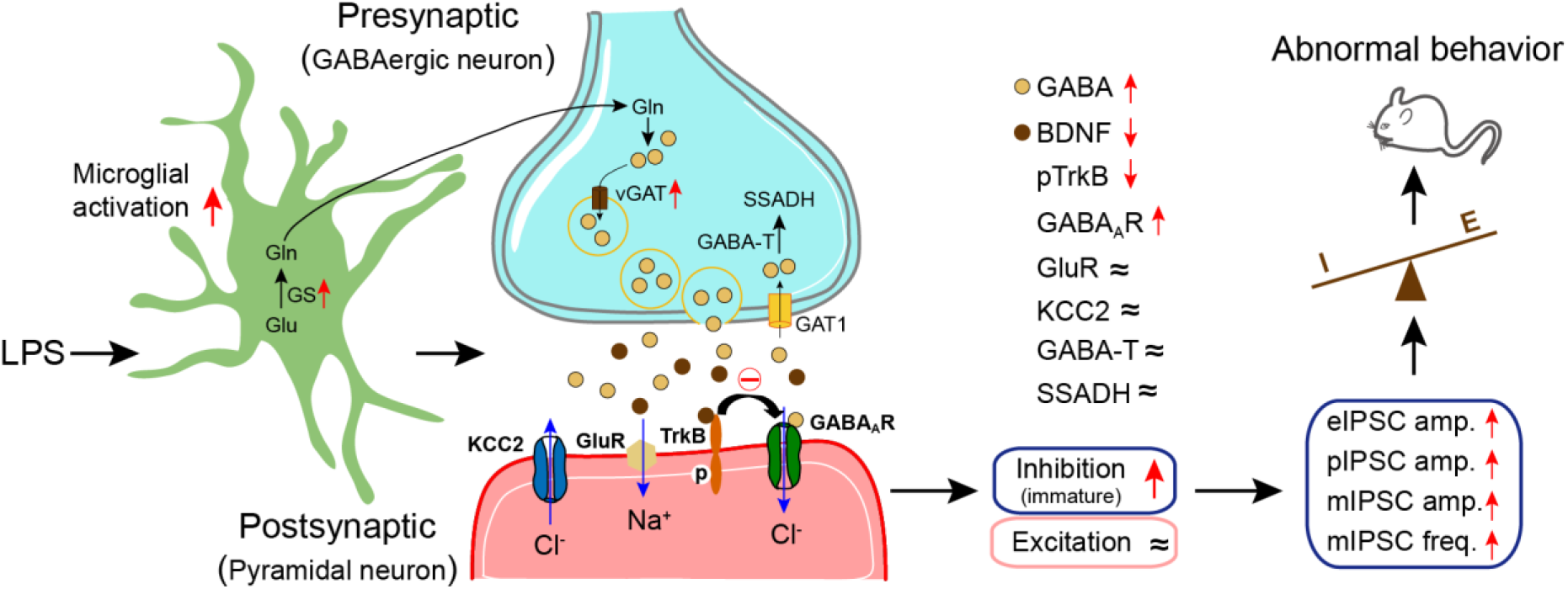
Diagram illustrating the mechanism underlying aberrant GABAergic synaptic transmission and subsequent behavior upon microglial activation. Microglia activation by LPS challenge results in enhanced amplitude of mIPSC due to upregulation of GABAAR subunits, α1 and β2, in glutamatergic neurons. Enhanced levels of GS and vGAT suggest increased GABA production and subsequent mIPSC frequency. In comparison, excitatory glutamate receptors (N1R, NR2B and GluR1), KCC2 and GABA degradation-related protein levels remain unchanged. BDNF is necessary for the maturation of GABAergic synapses and negatively regulates GABAAR-mediated IPSC. Down-regulation of BDNF together with reduced levels of pTrkB promote immature inhibitory response in the mPFC of 2h-LPS mice. Enhanced inhibition but unchanged excitation result in disturbance of E/I ratio in the mPFC, leading to aberrant mPFC govern behavior in mice.

Minocycline has long been used as an antibiotic for the treatment of various illnesses, demonstrating anti-inflammatory, immunomodulatory and neuroprotective effects **(Garrido-Mesa et al., 2013)**. Minocycline has recently shown promising results in the treatment of mental illness, such as bipolar disorder **(Savitz et al., 2018)**, depression **(Emadi-Kouchak et al., 2016)**, schizophrenia **(Zhang et al., 2018)** and cognitive impairment **(Chini et al., 2020)**. We observed that normalization of the levels of GABA_A_R subunits and vGAT by minocycline pretreatment, and the associated improvement in mIPSC amplitude and frequency, suggests that excessive microglial activation may have a direct influence on both pre- and postsynaptic activity at GABAergic synapses. Alternatively, retrograde postsynaptic signals, downstream of GABA_A_Rs, may also play a role in modulating presynaptic activities **(Duan et al., 2019)**. Blocking microglial activation with minocycline ameliorated abnormal behavior caused by LPS treatment as well, suggesting that altered microglial function due to acute neuroinflammation could trigger pathological behavior, which may be downstream of abnormal GABAergic synaptic function. Taken together, this study shows that acute neuroinflammation alters GABAergic synaptic transmission in the mPFC *via* activated microglia, and highlights the potential for the use of minocycline, which is generally considered an inexpensive and safe drug with a known side-effect profile in humans **(Garrido-Mesa et al., 2013)**, in the prevention and treatment of neuroinflammation-induced neuronal dysfunction and associated pathological behavior.

An electrophysiological parameter, half-width, indicates the maturity of postsynaptic currents, i.e., a broader half-width indicates less maturation, and vice versa **(Ali et al., 2000; Mitoh et al., 2008)**. GABA_A_RS, particularly the GABA_A_R α1 subunit, are critically involved in determining the kinetics of the inhibitory response and therefore the half-width of mIPSC. Upregulation of GABA_A_R α1 subunits usually reduces the half-width of fast inhibitory synaptic currents, a characteristic of mature inhibitory synapses **(Okada et al., 2000; Vicini et al., 2001)**. Strikingly, despite the upregulated levels of GABA_A_R α1 in LPS-challenged mPFC, we observed a significantly increased mIPSC half-width, which could be blocked by pretreatment with minocycline, in pyramidal neurons of 2 h-LPS mPFC. It has been clearly established that BDNF is necessary for the maturation of GABAergic synapses **(Fiorentino et al., 2009; Huang et al., 1999)**. Given that BDNF was significantly reduced in LPS-challenged mPFC, the present study suggests that excessive activation of microglia may promote an immature inhibitory response by downregulation of BDNF.

A consequence and marker of neuroinflammation due to microglial activation is the production of neurotoxic factors upon stimulation **(Calcia et al., 2016; Dantzer et al., 2008; Suzumura, 2013)**. In line with these findings, our RT-PCR analysis demonstrated significantly increased levels of mRNA for several proinflammatory cytokines, including TNF-α, interleukin 1β (IL-1β) and interleukin 6 (IL-6), in the mPFC of 2 h-LPS mice ***(Suppl. Fig. 3),*** confirming that neuroinflammation occurs as soon as 2 h after LPS insult. However, given that we observed an increase in GABAergic inhibition but unchanged glutamatergic excitation, it is unlikely that an increase in the levels of the above cytokines is a contributory factor. Indeed, numerous studies have shown that TNF-α decreases inhibitory synaptic strength by downregulation of levels of GABA_A_Rs, including the α1 and β2 subunits, due to increased internalization of GABA_A_RS **(Luscher et al., 2011; Olmos and Llado, 2014; Pribiag and Stellwagen, 2013; Stellwagen et al., 2005)** and decreased GABA release **(De Laurentiis et al., 2000)**. TNF-α is also known to increase excitatory synaptic strength **(De Laurentiis et al., 2000; Kawasaki et al., 2008; Olmos and Llado, 2014; Shim et al., 2018; Stellwagen et al., 2005; Zhang et al., 2013)**. Similarly, IL-1β and IL-6 also decrease GABAergic inhibition and increase glutamatergic excitation **(Atzori et al., 2012; Garcia-Oscos et al., 2012; Kawasaki et al., 2008; Wei et al., 2012)**; a contrasting result, showing that IL-6 significantly increased GABA release, was observed only under depolarizing conditions, i.e. with 40 mM K^+^ **(De Laurentiis et al., 2000)**, while the present study used physiological K^+^ levels, i.e. less than 3 mM. Together, this study suggests that upregulated GABAergic signaling may function as an early sensor of the infected brain. Thus, the involvement of microglial activation in the upregulation of GABAergic synaptic transmission in the mPFC, upon neuroinflammation, may require particular attention in treating inflammation-related brain dysfunction.

BDNF plays an important role in regulating synaptic transmission and plasticity in the CNS **(Carter et al., 2002; Patterson et al., 1996; Song et al., 2017)** and, in particular, in mediating microglia-synapse crosstalk **(Marinelli et al., 2019; Parkhurst et al., 2013)**. Microglia produce various neuron-protective factors, including BDNF **(Chen et al., 2014; Gomes et al., 2013; Parkhurst et al., 2013)**. Microglial displacement of inhibitory synapses decreases GABAergic inhibition, accompanied by upregulation of BDNF, in the motor cortex of 4-day LPS-treated mice, linking activated microglia with neuroprotection **(Chen et al., 2014)**. BDNF decreases the cell surface expression of GABA_A_Rs-gephyrin complex **(Mou et al., 2013)** and suppresses GABAergic synaptic transmission via activating endocytosis and repression of GABA_A_RS **(Riffault et al., 2014)**. Therefore, these findings implicate a negative correlation between BDNF and GABAergic inhibition. Strikingly, we demonstrate here that microglial activation downregulates levels of BDNF and this is associated with enhanced GABAergic synaptic transmission of pyramidal neurons in the mPFC. Our result is consistent with above and other findings showing that neurotrophies, including BDNF, reduce inhibitory synaptic transmission **(Brunig et al., 2001; Frerking et al., 1998; Kim et al., 1994; Tanaka et al., 1997)**. The consensus in the literature is that BDNF controls a wide spectrum of processes, albeit sometimes with contradictory effects, which can be explained either by its synthesis pattern, or different active isoforms binding to various types of receptor, or the triggering of different signaling pathways **(Kowianski et al., 2018; Martinowich et al., 2007; Wardle and Poo, 2003)**. The present study again underlines the notion that it is important to correlate BDNF functions with the stage of brain development, the particular brain area, and the various cellular components and signal-transduction mechanisms involved in the molecular physiology of both healthy and pathological states **(Kowianski et al., 2018; Martinowich et al., 2007)**.

Furthermore, we have shown that upregulating levels of BDNF, by blocking microglial activation with minocycline or by exogenous application of BDNF, ameliorates abnormal mIPSCs in pyramidal neurons of the mPFC. Indeed, our results implicate BDNF in a novel molecular pathway that mediates GABAergic synaptic dysfunction due to microglial activation. Upon binding BDNF, monomeric TrkB receptors dimerize, autophosphorylate, and as a result activate signaling pathways **(Chao, 2003)**. It is worth noting that LPS only reduces levels of pTrkB, in addition to BDNF, while leaving overall levels of TrkB unchanged. Moreover, minocycline normalizes levels of BDNF and pTrkB without affecting TrkB levels. This suggests that minocycline might increase the ability of TrkB to bind its ligand, BDNF **(Carter et al., 1995)**, and thereby enhance the activation of the downstream molecular pathways necessary for maintaining normal GABAergic function in the mPFC of 2 h-LPS mice.

A recent study showed that upregulation of glutamatergic synaptic transmission occurs in pyramidal cells of hippocampal CA1 within 2 h of the systemic inflammation resulting from LPS challenge **(Duan et al., 2018)**. In contrast, we only observed enhanced GABAergic synaptic transmission in the mPFC pyramidal cells of inhibitory-excitatory synapses, leaving the excitatory response unchanged. However, Duan et al. used P14 mice, in which LPS injection does not induce significant levels of activated microglia in the CA1 of hippocampus **(Duan et al., 2018)**. It is known that microglia exhibit brain region-dependent heterogeneity in their core gene profile **(De Biase et al., 2017; Grabert et al., 2016)**, and differential response of hippocampal and PFC oscillations to systemic LPS treatment has been reported in a study of local field potential which suggests a region-specific neuronal activity in LPS-induced acute inflammation **(Mamad et al., 2018)**. Our study, conducted in 1-2 month-old mice, supports the notion that microglia in different brain regions, or across different developmental or pathogenic phases, may rely on different signaling mechanisms to perform various functions **(Cantaut-Belarif et al., 2017; Miller-Rhodes, 2018)**.

Finally, this study offers a new perspective on the postsynaptic cell-type-specific regulation of GABAergic synaptic transmission in the mPFC during acute neuroinflammation. Our results suggest that increased GABAergic, but not glutamatergic, synaptic transmission of the mPFC pyramidal neurons is one of the major changes associated with microglial activation and abnormal behavior during the early phase of neuroinflammation, which advances our knowledge of microglia-synapse interaction. Giving that the GABAergic system has great potential to inhibit inflammatory responses **(Bhat et al., 2010; Jin et al., 2013; Wu et al., 2017)**, the present findings highlight the potential of components of the GABAergic system as new therapeutic targets for inflammatory diseases. Moreover, microglial regulation of GABAergic synaptic transmission may not be restricted to disease states, and the mechanisms that we have deciphered here may be relevant to healthy physiological situations, which deserves further exploration in future studies.

## Materials and Methods

### Mice and Treatments

All experimental procedures were approved by the Guangzhou University Animal Care and Use Committee, according to the Guidelines for Animal Care established by the National Institute of Health. C57BL/6J mice were obtained from the Guangdong Medical Laboratory Animal Center. GAD67^+/GFP^ mice **(Tamamaki et al., 2003)** were a gift from Dr. Yuqiang Ding at the School of Medicine of Tongji University. Mice used in this study were 1-2 month-old male and female mice. Neuroinflammation was induced by single-dose interperitoneal (IP) injection of LPS (Sigma Aldrich, 0.5 mg/kg). LPS was diluted in phosphate-buffered saline (PBS) (0.01 M, pH 7.4). PBS (indicated as Ctrl group in the text) or LPS was injected 2 h prior to experimental measurements. For minocycline treatment, mice were pretreated with either minocycline (50 mg/kg, IP, Sigma Aldrich) or PBS once daily for three consecutive days, followed by a single IP injection of LPS (0.5 mg/kg) or PBS on the third day.

### Immunofluorescence

Mice were anesthetized by IP injection of 20% urethane (0.01 ml/g) and perfused transcardially with 0.9% saline and 4% paraformaldehyde (PFA) in PBS (0.01 M, pH 7.4). Following perfusion, the brains were dissected and post-fixed overnight in 4% PFA followed by a sucrose solution series (10%, 20% and 30%) at 4°C for cryoprotection. Serial coronal/sagittal sections of tissue were obtained using a Leica CM30505 with a thickness of 30 μm. Free-floating sections were rinsed in PBS three times for 5 min and processed for antigen retrieval by boiling in 10 mM citrate buffer (pH 6.0) for 5 min. The sections were incubated in 3% H_2_O_2_ in PBS for 15 min at room temperature (RT) to quench endogenous peroxidases, washed three times for 5 min and subsequently incubated in 0.3% Triton X-100 in PBS for 2 h. To block nonspecific binding, sections were incubated in 3% bovine serum albumin (BSA) for 30 min. Sections were incubated with antibodies against Iba1 (1:1000) overnight at 4°C in 1% BSA/PBS. After three washes with PBS, secondary antibody (Invitrogen) application was performed at RT for 2 h followed by three additional washes with PBS for 10 min. Sections were then washed and mounted on gelatin-coated slides, and imaged on a fluorescence microscope.

### Western Blotting

Mouse brains were homogenized with micro tissue grinders (Kimble) in 2 mL tubes, using lysis buffer containing 50 mM Tris pH 7.5, 150 mM NaCl, 5 mM EDTA pH 8.0, 1% SDS, and protease inhibitor (Complete Mini; Roche). After 1 min homogenization, cellular debris was removed by centrifugation at 14000×g for 10 min at 4°C, and supernatant was collected for denaturation at 75°C for 20 min. Tissue lysates were subjected to SDS-PAGE (Bio-Rad) and transferred to nitrocellulose membranes. The membranes were blocked for 1 h using 5% non-fat dry milk in Tris-buffered saline (TBS) containing 0.5% Tween-20 (TBST). Membranes were then probed with primary antibodies against NMDAR1 (NR1, Abcam, ab109182), glutamate receptor 1 (GluR1, AMPAR subtype, Abcam, ab31232), NMDAR2B (NR2B, Abcam, ab65783), GS (Abcam, ab49873), vGAT (Abcam, ab101999), GABA_A_R subunits, α1 (Abcam. ab86320), α5 (Abcam, ab175195), β2 (Abcam, ab156000), BDNF (Abcam, ab203573), TrkB (Genetex, GTX13372), pTrkB (Tyr816) (Sigma, ABN1381), KCC2 (Millipore, 07-432), GABA-T (Abcam, ab152134), SSADH (Abcam, ab129017). Anti γ-tubulin (Abcam, ab87328) or anti-β-actin antibody (Abcam, ab20272) was used as loading control. After three washes with TBST, HRP-labeled secondary antibody (CWS) application was performed at RT for 1 h using 5% milk in TBST followed by three additional washes with TBST. Bands were visualized using the Immobilon Western ECL system (Millipore) and analyzed with Gel Pro Analysis software **(Yang et al., 2009)**.

### Quantitative Real-Time PCR

Total RNA was extracted from dissected mPFC tissues with Trizol reagent (Vazyme Biotech, Nanjing following the manufacturer’s protocol. RNA concentrations were measured by the absorbance at 260 nm (NanoDrop 2000 spectrophotometer, Thermo Fisher Scientific, Wilmington, DE) and 1 μg total RNA was used for the synthesis of first-strand cDNA (PrimeScript™ RT Master Mix, TakaraBio Inc., Shiga, Japan). Quantitative real-time PCR (RT-PCR) was conducted on a CFX96 Touch RT-PCR Detection System (Bio-Rad, Hercules, CA) with SYBR Premix Ex Taq TM II (Takara Bio). The specificity of amplicons was examined by melting curve analysis and agarose gel electrophoresis. Relative expression levels of target genes were determined by the 2^−ΔΔCt^ method **(Livak and Schmittgen, 2001)** with glyceraldehyde 3-phosphate dehydrogenase (GAPDH) as the endogenous reference gene. Primer sequences were shown in ***Table S1***.

### Whole-Cell Patch Clamp Recording

mPFC slices (350 μm) were cut using a vibratome (Leica VT1000S) in ice-cold cutting solution containing the following: 119 mM NaCl, 2.5 mM KCl, 2.5 mM CaCl_2_, 1.3 mM MgSO4, 1 mM NaH2PO4, 11 mM D-glucose, 26.2 mM NaHCO3. Pipettes were filled with an internal solution of 136.5 mM K-gluconic acid, 17.5 mM KCl, 9 mM NaCl, 1 mM MgCl_2_, 0.2 mM EGTA, 10 mM HEPES (pH 7.3) containing gramicidin A 10 mg/ml (Sigma; diluted from a stock solution of 10 mg/ml in DMSO). The recording aCSF contained 140 mM NaCl, 4.7 mM KCl, 2.5 mM CaCl_2_, 1.2 mM MgCl_2_, 11 mM D-glucose, 10 mM HEPES (pH 7.3). Both incubating and recording solutions were bubbled with 95% O_2_ and 5% CO_2_ throughout the experiment. mPFCs were dissected, weighed, flash-frozen in liquid nitrogen and then stored in a −80°C freezer. To make the mPFC homogenate for puffing experiments, two freeze–thaw cycles were performed to break the cell membrane before adding 20 μl bath solution per milligram tissue into the tube, followed by 1 min sonication on ice. The homogenate was centrifuged for 5 min at 5000×g, and supernatant of PBS or LPS mPFC was then aliquoted and stored at −20°C. The stored mPFC supernatant was further diluted using aCSF (1:100) for puffing experiments. Puffs were of 200 ms duration at 50 psi and were delivered using a Picospritzer (Parker Instrumentation). Voltage was held at −60 mV and 0 mV, respectively, for mEPSC and mIPSC recordings **(Chen et al., 2017; Liang et al., 2015)**, the aCSF was supplemented with 1 mM tetrodotoxin (TTX), and pipettes were filled with an internal solution of 100 mM Cs gluconate, 5 mM CsCl, 10 mM HEPES, 2 mM MgCl_2_, 1 mM CaCl_2_, 11 mM BAPTA, 4 mM ATP, 0.4 mM GTP. Data were filtered during acquisition with a low pass filter set at 2 kHz using pClamp10 (Molecular Devices). Data analysis was performed offline with Clampfit 10.2 (Molecular Devices).

To evaluate the effect of BDNF on LPS-induced abnormal mIPSC, brain slices were incubated in aCSF with LPS at final dose of 50 ng/ml, 100 ng/ml or 200 ng/ml, respectively. Slices were preincubated with BDNF (Bio-Techne, 248-BDB) at 20 ng/ml or 50 ng/ml, 20 min prior to LPS application.

### Open Field Test (OFT)

The OFT was performed as previously described **(Misrani et al., 2019)**. Briefly, after adaptation for 1 h in the experiment room, mice were placed directly in the center of the open field. The movement of each animal in the area was video-recorded during the 5 min testing session. After 5 min, the animal was removed and returned to the home cage, and the open field area was cleaned with 75% ethanol to prevent olfactory cues from affecting the behavior of subsequently tested mice. Exploration was defined as the time spent in the center of the nine squares, and overall activity was defined as the number of squares crossed during the testing session. The longer the time animals spent in the center of the nine squares, the lower their anxiety levels **(Kraeuter et al., 2019; Seibenhener and Wooten, 2015)**.

### Elevated Plus Maze (EPM)

Anxiety-like behavior was detected by the EPM test as previously described **(Chen et al., 2017)** with little modification. The EPM apparatus consisted of two open arms, two closed arms, and a common central platform. Briefly, after adaptation for 1 h in the experiment room, mice were placed directly on the center platform of the EPM apparatus facing the open arm. The total number of arm entries and time spent in the open arms were video recorded for 5 min; mice were then removed and returned to their home cages. Following each test, the EPM area was cleaned with 75% ethanol.

### Tail Suspension Test (TST)

The total duration of immobility in the TST was measured according to the method described **(Leger et al., 2015)**. Briefly, mice were suspended by the tail with a paper clip attached with adhesive tape about 1 cm from the end of the tail. Immobility time was recorded during a 6 min period by an experienced observer blind to the experimental groups. Mice were considered immobile only when they hung passively and completely motionless.

### Sucrose Preference Test (SPT)

The SPT consisted of a two-bottle choice paradigm **(Scheggi et al., 2018; Yan et al., 2018)**. Mice were first habituated to two bottles of 1% sucrose solution or tap water for three days. The bottle positions were swapped once every 24 h to establish baseline preference. For the next three days, the mice were given access to two pre-weighed bottles containing 100 ml water and 100 ml 1% sucrose solution, respectively. The position of the two bottles was swapped every 24 h in order to discriminate between a positional preference and sucrose preference. Bottles were weighed every day to measure sucrose consumption.

### Statistics

Statistical analysis was performed using OriginPro 2017 software with either Student’s *t*-test for two-group comparisons, one- or two-way ANOVA with a Bonferroni’s *post hoc* test or Tukey’s multiple comparisons. *P* < 0.05 was considered statistically significant.

## Acknowledgements

This work was supported by grants from the National Natural Science Foundation of China (31771219, 31970915, 31871170, 81804197), Guangdong Natural Science Foundation for Major Cultivation Project (2018B030336001).

## Author Contributions

Binliang Tang, Conceptualization, Recorded and analyzed electrophysiology data, conducted behavior test, Writing – original draft; Jinxiang Jiang, Western blotting and data analysis, Recorded and analyzed electrophysiology data, Statistics, Writing – original draft; Lei Wang, Iba1 staining and real-time PCR, Writing – original draft; Afzal Misrani, Data acquisition, Methodology; Qingwei Huo, Visualization, Statistics; Yuanyuan Han, GFAP staining; Cheng Long, Supervision, Funding acquisition; Li Yang, Supervision, Funding acquisition, Project administration, Writing – original draft, Writing – review and editing.

## Notes

**Conflict of interest:** No conflict of interest

### Competing Interest Statement

The authors have declared no competing interest.

