## Supplemental figures and table for "Microglial activation results in neuron-type-specific increase in mPFC GABAergic transmission and abnormal behavior in mice"

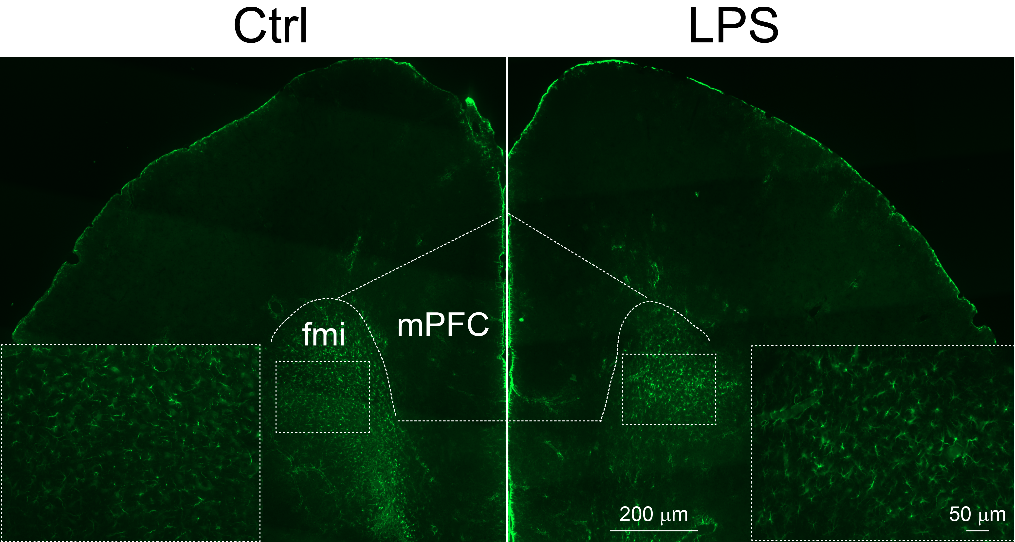

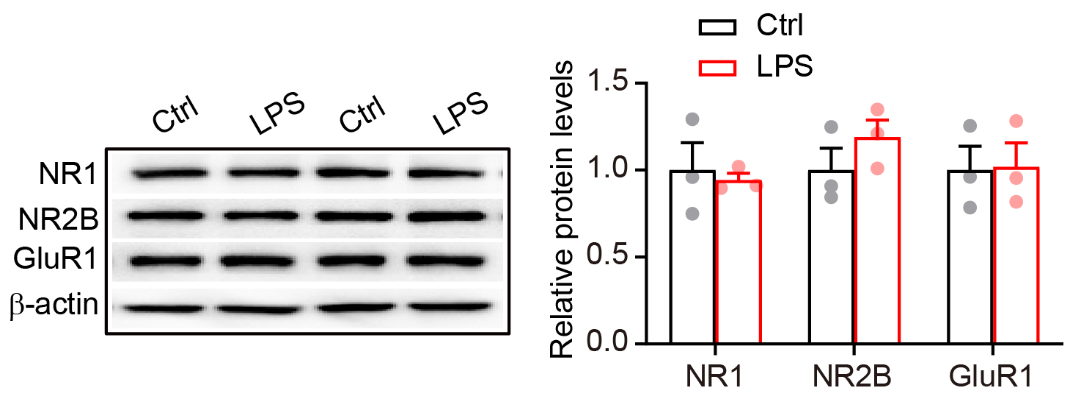
**Figure S1**. Unchanged levels of excitatory receptors in the mPFC of LPS mice. **(A)** Representative western blotting bands and **(B)** quantification of levels of NR1, NR2B and GluR1 in the mPFC (n=3 for each group). Data are presented as means ± SEM. Student’s *t*-test.

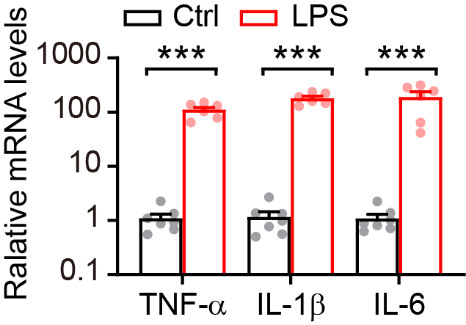
**Figure S2**. Immunofluorescent staining of GFAP on the mPFC sections showing similar level of astrocyte activation in Ctrl and LPS mice.

**Figure S3**. Real-time PCR analysis showing elevated mRNA levels of pro-inflammatory cytokines, TNF-α, IL-1β and IL-6, in the LPS-treated mPFC (Ctrl, n=3; LPS, n=4). Data are presented as means ± SEM. Student’s *t*-test, ****P*<0.001.

**Table S1.** Primer sequences of real time-PCR

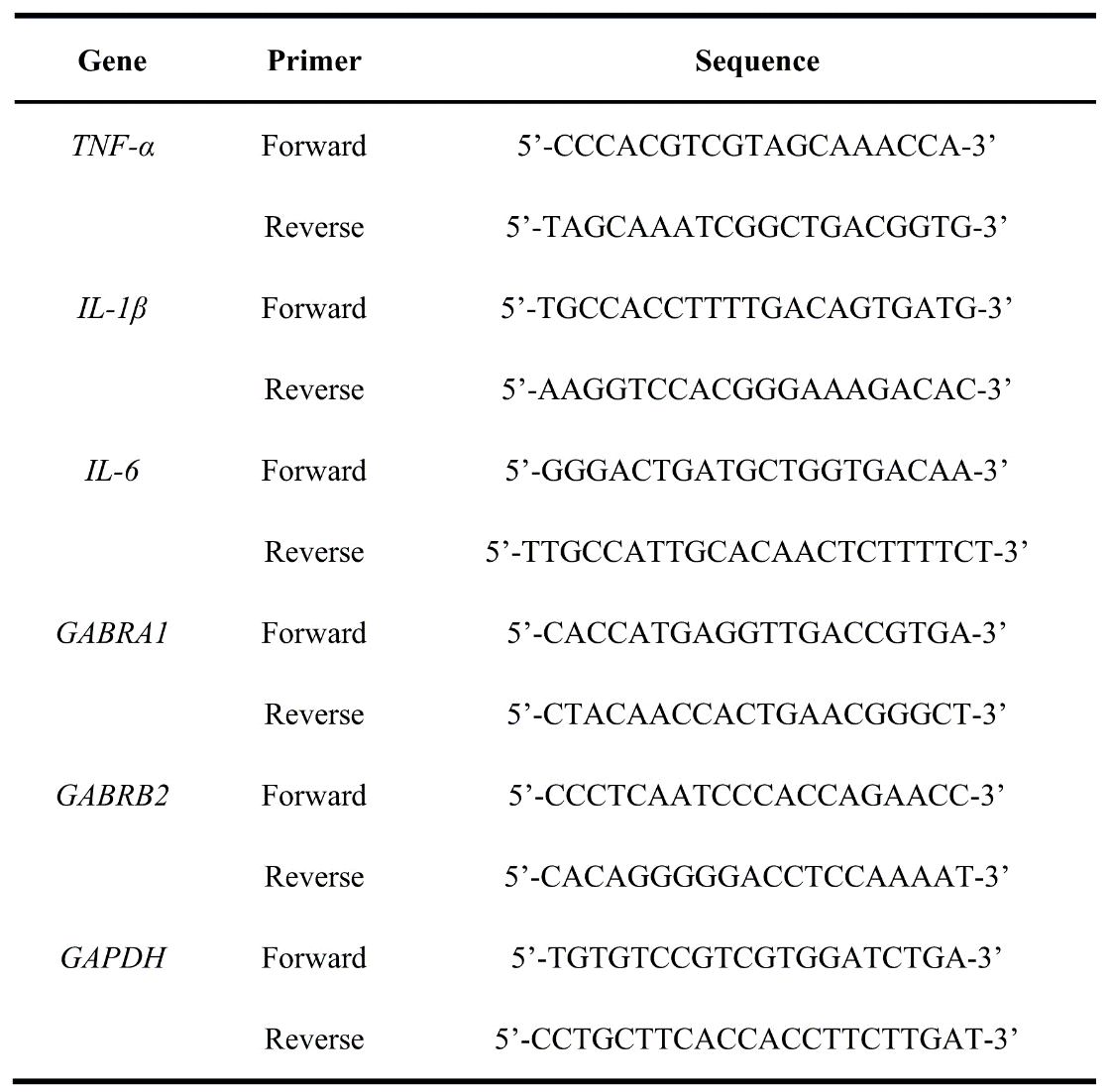
